# Diverging maternal and infant cord antibody functions from SARS-CoV-2 infection and vaccination in pregnancy

**DOI:** 10.1101/2023.05.01.538955

**Authors:** Emily H. Adhikari, Pei Lu, Ye jin Kang, Ann R. McDonald, Jessica E. Pruszynski, Timothy A. Bates, Savannah K. McBride, Mila Trank-Greene, Fikadu G. Tafesse, Lenette L. Lu

**Affiliations:** Division of Maternal-Fetal Medicine and Department of Obstetrics and Gynecology, UTSW Medical Center, Dallas, TX; Parkland Health, Dallas TX; Division of Infectious Diseases and Geographic Medicine and Department of Internal Medicine, UTSW Medical Center, Dallas, TX; Department of Microbiology and Immunology, Oregon Health and Science University, Portland, OR; Department of Immunology, UTSW Medical Center, Dallas, TX

## Abstract

Immunization in pregnancy is a critical tool that can be leveraged to protect the infant with an immature immune system but how vaccine-induced antibodies transfer to the placenta and protect the maternal-fetal dyad remains unclear. Here, we compare matched maternal-infant cord blood from individuals who in pregnancy received mRNA COVID-19 vaccine, were infected by SARS-CoV-2, or had the combination of these two immune exposures. We find that some but not all antibody neutralizing activities and Fc effector functions are enriched with vaccination compared to infection. Preferential transport to the fetus of Fc functions and not neutralization is observed. Immunization compared to infection enriches IgG1-mediated antibody functions with changes in antibody post-translational sialylation and fucosylation that impact fetal more than maternal antibody functional potency. Thus, vaccine enhanced antibody functional magnitude, potency and breadth in the fetus are driven more by antibody glycosylation and Fc effector functions compared to maternal responses, highlighting prenatal opportunities to safeguard newborns as SARS-CoV-2 becomes endemic.

**One Sentence Summary:** SARS-CoV-2 vaccination in pregnancy induces diverging maternal and infant cord antibody functions

## INTRODUCTION

With over 140 million women giving birth every year, the newborn immature immune system and maternal immune adaptation to pregnancy represent widespread challenges to surviving COVID-19. Infants > 6 months old have higher rates of hospitalization compared to those older for SARS-CoV-2 infection (1, 2). Similarly, pregnant individuals are more likely to need intensive care unit and ventilatory support compared to the general population (3, 4), and have increased risk of obstetrical and fetal complications (5, 6). Beyond the short-term consequences, a growing number of long-term multi-system sequelae for adults are being recognized (7). Moreover, emerging data suggest that even without vertical transmission there may be infant neurodevelopmental impacts from prenatal exposure to SARS-CoV-2 yet to be fully understood (8-10). Thus, immunizations could be critical to safeguarding maternal, fetal and newborn health as SARS-CoV-2 becomes endemic and public health precautions wane but how they can be harnessed to provide optimal protection are less clear.

Before vaccine availability, COVID-19 contributed to ∼25% of maternal deaths (11, 12) and was the number one cause of infection-related deaths in children (13). Recent studies show that as in the general population, monovalent mRNA immunization in pregnancy decreases the risk of maternal and infant SARS-CoV-2 infection, disease, and mortality (14-19). Unraveling the maternal responses to the different immune exposures of infection and vaccination during pregnancy and what transfers into the placenta can guide vaccine design and implementation strategies to enhance protection of the maternal-fetal dyad.

Multiple human and animal studies show that leveraging a breadth of immune responses to differentially target several steps in viral infection and pathogenesis is likely necessary to provide durable protection (20-25). After mRNA vaccination, the primary form of immunity transferred to the fetus is antibodies, specifically IgG. If we understand the spectrum of IgG-mediated functions in the fetus then we can design ways to leverage maternal immunity to protect the infant.

SARS-CoV-2 mRNA vaccines and infection generate IgG with neutralizing and antibody Fc effector functions in the non-pregnant (26-29) and pregnant populations (30-35). With Fc-Fc receptor engagement on innate and adaptive immune cells (36, 37), induced effector functions can prevent disease even after infection has occurred by eliminating infected cells and blocking spread (25, 38-44). Diversity of the Fc domain through differential subclass and post-translational glycosylation modulates binding to Fc receptors and the spectrum of effector functions (45-52). Moreover, antibody functional potency, breadth and coordination between neutralizing and Fc effector responses likely contribute to protection (26, 53-57). Some data demonstrate that maternal immune adaptation to pregnancy alters antibody subclass and glycosylation (30, 58-65), but the implications for antibody functions and what exists for the fetus are only just beginning to be appreciated (66-70).

To understand how SARS-CoV-2 mRNA vaccination in pregnancy impacts the newborn, we collected paired maternal-infant cord blood samples at delivery. We evaluated neutralization against live SARS-CoV-2 and the Fc effector functions of natural killer cell activation that leads to antibody-dependent cellular cytotoxicity, antibody-dependent monocyte phagocytosis, antibody-dependent complement deposition, and Fcγ receptor binding specific for the Spike glycoprotein receptor binding domain (RBD). We determined relative levels of RBD-specific antibody subclass, isotype and post-translational glycosylation to assess how these different features contribute to function. The data show that compared to SARS-CoV-2 infection, vaccination in pregnancy enhances some but not all neutralizing and Fc effector functions, with preferential transport of Fc functions and not neutralization. All functions are primarily driven by IgG1 which is enhanced by vaccination. However, vaccination in pregnancy changes glycosylation of cord and not maternal RBD IgG and the impact of glycosylation on antibody functional potency is observed more in cord compared to maternal responses. Thus, while vaccination compared to infection in pregnancy boosts antibody functions, the maternal and fetal paths begin to diverge.

## RESULTS

### Study subjects

We collected paired maternal-umbilical cord blood at deliveries from individuals who during pregnancy received mRNA vaccination targeting WA1 RBD (n=19 vaccine only) and or infected with SARS-CoV-2 (n=22 infection only and n=28 both vaccine and infection) at Parkland Health, the Dallas County’s public hospital in Texas (Table and Supplemental Table 1). Clinical groups were defined by clinical history, documented vaccination in pregnancy, SARS-CoV-2 nasal swab PCR and SARS-CoV-2 nucleocapsid IgG (Supplemental Figure 1). Of the 47 vaccinated, four received mRNA-1273 and 43 BNT162b2 with no significant difference between the two (Supplemental Table 2). The mean age of individuals with only infection was lower compared to those who received vaccination in pregnancy (p=0.005). Body mass index (BMI), gestational age at delivery (75% full term), infant sex and an additional 21 clinical outcomes were not significantly different (Table and Supplemental Table 1). Of the infected, 20% were asymptomatic, 40% mild, 10% moderate, 14% severe and 16% critical. Of the vaccinated, the majority received two doses (68%) before delivery with their last dose in the third trimester (Supplemental Figure 1). With 94% of participants Hispanic, this study population represents a subset of the 11,170 deliveries at Parkland Health in 2021 and a patient population from social-economic communities disproportionately impacted by SARS-CoV-2 (Supplemental Table 3) (71-73).

**Table.** Clinical characteristics of study patients

| **Characteristic** | **Infection**<br>**n = 22** | **Vaccine**<br>**n = 19** | **Vaccine and**<br>**infection**<br>**n = 28** | **P value** |
| --- | --- | --- | --- | --- |
| Age, years | 28.3 ± 6.5 | 32.4 ± 5.8 | 34.4 ± 6.7 | 0.005 |
| Race/ethnicity |  |  |  | 0.151 |
| Hispanic | 20 (91) | 17 (90) | 28 (100) |  |
| Black, non-Hispanic | 2 (9) | 1 (5) | 0 (0) |  |
| White, non-Hispanic | 0 (0) | 1 (5) | 0 (0) |  |
| Other |  |  |  |  |
| Maternal BMI at first visit, kg/m<sup>2</sup> | 32 (26-36) | 30 (27-33) | 34 (30-39) | 0.147 |
| Male infant sex | 12 (55) | 12 (63) | 15 (54) | 0.789 |
| EGA at delivery, weeks | 38 (36-38) | 38 (38-39) | 37 (37-38) | 0.300 |

Data shown as n (%), mean ± standard deviation (SD), or median (Q1-Q3) as appropriate. BMI= body mass index, EGA= estimated gestational age

### Infant-cord SARS-CoV-2 neutralizing and Fc effector functions

Infants of individuals with monovalent COVID-19 mRNA vaccination in pregnancy have decreased risk of SARS-CoV-2 infection, hospitalization, and mortality up to 6 months after birth (15, 19). This is irrespective of whether SARS-CoV-2 infection occurred. To begin to investigate the antibody functions induced by vaccination, we measured neutralizing activity that prevents viral entry and infection (74-77). Through focus reduction neutralization tests (FRNT) using live SARS-CoV-2 (WA1/2020) and variants Delta (B.1.617.2) and Omicron (BA.2) (Supplemental Figure 2), we observed higher cord neutralizing activities against WA1 in those with mRNA vaccination compared to SARS-CoV-2 infection and non-significant changes against Delta and Omicron (Figure 1A, F). Subanalyses showed that while neutralizing activity from the combination of vaccine and infection compared to vaccine alone trended higher, this was not statistically significant (Figure 1F). Thus, vaccination compared to infection alone in pregnancy induces higher cord neutralizing activity for protection against subsequent challenges by the same viral strain.

**Figure 1:**
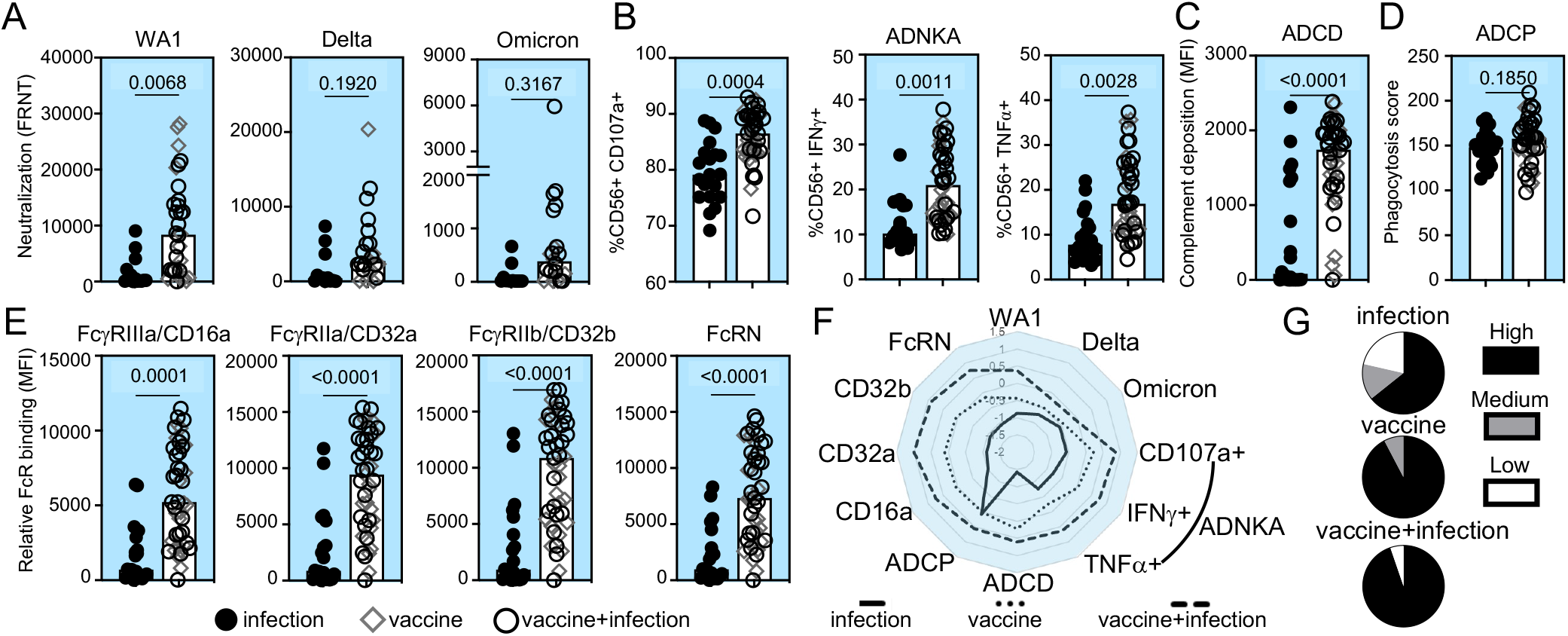
A subset of infant cord SARS-CoV-2 neutralizing and RBD Fc effector antibody s is enhanced with vaccination compared to infection in pregnancy. The medians (bars) sample (**A**) neutralization (FRNT50) against SARS-CoV-2 WA1 (infection n=14, vaccine accine+infection n=19), Delta and Omicron viruses (infection n=12, vaccine n=8, infection n=14), (**B**) RBD antibody-dependent natural killer cell activation (ADNKA) by, IFNγ, and TNFα, (**C**) RBD antibody-dependent complement deposition (ADCD), (**D**) ibody-dependent cellular phagocytosis (ADCP), and (**E**) relative binding of RBD-specific es to FcγRIIIa/CD16a, FcγRIIa/CD32a, FcγRIIb/CD32b and FcRN are shown. For B-E, sizes are infection n=20, vaccine n=18, vaccine+infection n=27. P-values for A-E are for maternal age and body mass index using linear regression. (**F**) The magnitude of ctions are summarized in the radar plot. Each line represents the median Z-scored data clinical group. (**G**) The proportion of detectable functions was used to categorize ls as a high, medium or low responder. The percentages of each type of responder ch clinical group depict the polyfunctional antibody breadth.

Decay studies show that antibody Fc effector functions are more durable than neutralization (23, 29, 78) and data from animal models show that Fc-Fc receptor engagement impacts viral load and disease (25, 38-42, 44, 57). We examined RBD-specific Fc effector functions induced by vaccination and SARS-CoV-2 infection (26-28, 30, 32, 43, 46): antibody-dependent natural killer cell activation (ADNKA) which leads to antibody-dependent cellular cytotoxicity (ADCC) (Figure 1B), antibody-dependent complement deposition (ADCD) (Figure 1C) and antibody-dependent cellular phagocytosis (ADCP) (Figure 1D). We found that vaccination in pregnancy compared to infection alone is linked to higher RBD ADNKA including CD107a degranulation and intracellular IFNγ and TNFα production (Figure 1B) and ADCD (Figure 1C), not ADCP (Figure 1D). Antibody Fc domain engagement of low affinity Fc receptors is the first step in the signaling and initiation of effector functions while the high affinity FcRN is responsible for transport across the placenta and recycling (37, 79-81). We found that binding to the low affinity activating FcγRIIIa/CD16a and FcγRIIa/CD32a, the inhibitory FcγRIIb/CD32b and high affinity FcRN were elevated after vaccination compared to infection in pregnancy (Figure 1E). Again, the combination of vaccine and infection compared to vaccine alone was not statistically different (Figure 1F). Thus, the magnitude of some but not all cord RBD Fc effector functions and Fc receptor binding are enhanced with vaccination.

Sex (82, 83), infant prematurity (69), trimester of vaccination (31, 33), mRNA vaccine platform (27, 33) and disease severity (45, 46, 84) impact antibody responses. Though limited by power, no significant differences were observed in these data. Consistent with findings in the general population (85), there were no significant differences with respect to the order of vaccination and infection in the combination group (Supplemental Figure 1).

In addition to magnitude, greater polyfunctional antibody breadth is associated with increased protection (26). For each cord sample, we categorized the proportion of detectable SARS-CoV-2 neutralizing and Fc effector responses as high (>90%), medium (80-90%) or low (<80%) responder (Supplemental Figure3). We observed more high responders with greater functional breadth in the vaccinated compared to infected (Figure 1G).

### Maternal SARS-CoV-2 neutralizing and Fc effector functions

To understand how infant cord antibody responses are shaped, we next measured neutralizing activities and Fc effector functions in paired maternal samples obtained at delivery. Like cord samples, maternal blood from those vaccinated compared to infected enhanced the magnitude of neutralization against SARS-CoV-2 WA1 and not Delta or Omicron (Figure 2A). RBD ADNKA (Figure 2B), ADCD (Figure 2C), and not ADCP (Figure 2D) were increased. Relative binding of RBD IgG to FcγRIIIa/CD16a, FcγRIIa/CD32a, FcγRIIb/CD32b and FcRN were higher (Figure 2E). There was a non-statistically significant trend towards increased antibody functions with combination vaccine and infection compared to vaccine alone (Figure 2F) and greater polyfunctional breadth with vaccination (Figure 2G). Thus, vaccination in pregnancy enhances the magnitude and breadth of neutralizing and Fc effector functions across the maternal-fetal dyad.

**Figure 2:**
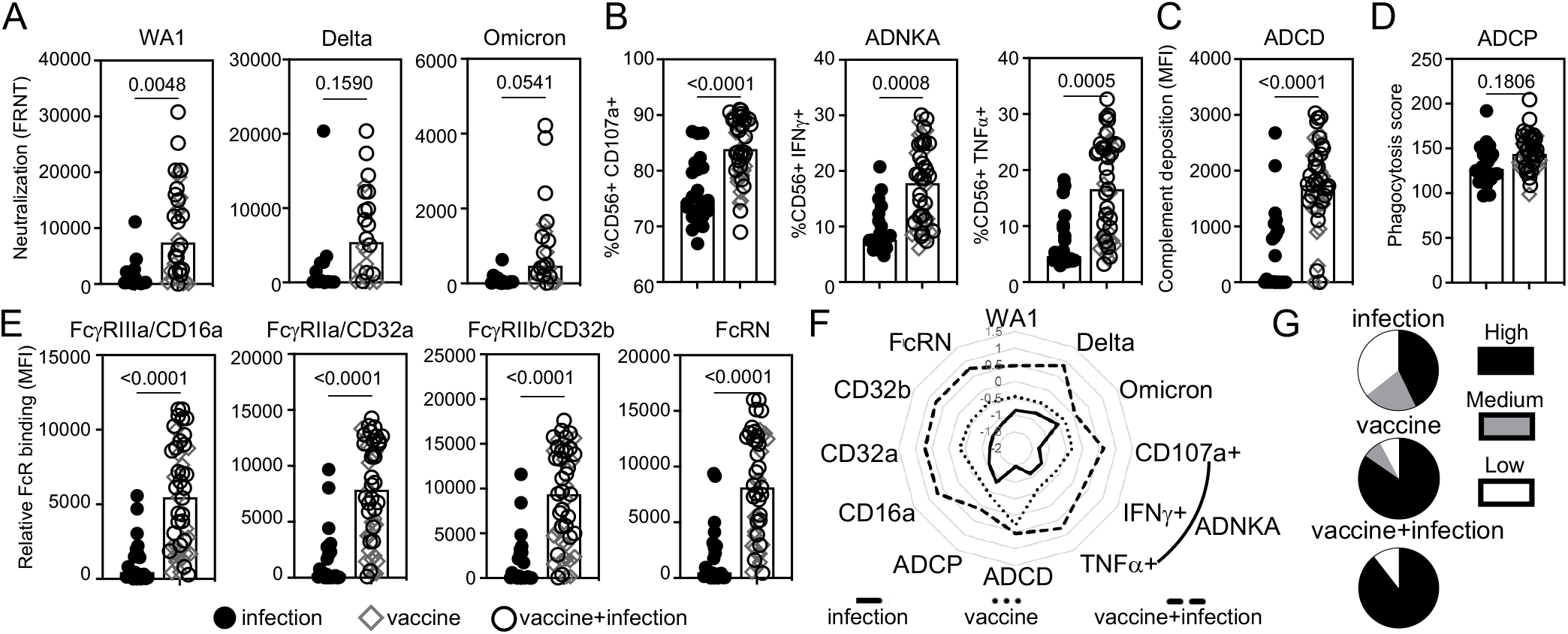
A subset of maternal SARS-CoV-2 neutralizing and RBD Fc effector antibody functions ced with vaccination compared to infection in pregnancy. The medians (bars) of the l pair of the cord samples in Figure 1 in (**A**) neutralization (FRNT50) against SARS-CoV-(infection n=14, vaccine n=13, vaccine+infection n=19), Delta and Omicron viruses n n=12, vaccine n=8, vaccine+infection n=14), (**B**) RBD antibody-dependent natural killer ation (ADNKA) as measured by CD107a, IFNγ, and TNFα, (**C**) RBD antibody-dependent ent deposition (ADCD), (**D**) RBD antibody-dependent cellular phagocytosis (ADCP), relative binding of RBD-specific antibodies to FcγRIIIa/CD16a, FcγRIIa/CD32a, CD32b and FcRN. For B-E, sample sizes are infection n=22, vaccine n=19, infection n=28. P-values for A-E are adjusted for maternal age and body mass index ear regression. (**F**) The magnitude of maternal functions are summarized in the radar h line represents the median Z-scored data for each clinical group. (**G**) The proportion of le functions was used to categorize individuals as a high, medium or low responder. The ges of each type of responder within each clinical group depict the polyfunctional breadth.

### Maternal-fetal transfer of antibody functions

The maternal response to an immune exposure and transfer of that response across the placenta determine fetal antibody functions. In examining transfer, we found that neutralizing activities did not differ between paired maternal and cord samples irrespective of immune exposure (Figure 3A). However, levels of RBD Fc effector functions of ADNKA were lower in maternal compared to cord blood (Figure 3B). This was not observed for RBD ADCD (Figure 3C) but was for ADCP (Figure 3D). Consistent with the low affinity activating and inhibitory FcγRs modulating ADNKA and ADCP, relative binding for FcγRIIIa/CD16a, FcγRIIa/CD32a and FcγRIIb/CD32b were all lower in maternal compared to cord blood (Figure 3E). In contrast, no difference was observed for the high affinity FcRN that mediates IgG transport across the placenta (Figure 3E). These data show preferential transfer of a subset of Fcγ effector functions compared to viral neutralization and C1q mediated C3 complement deposition that leads to ADCD. In the case of Fcγ receptor binding the transfer ratio was inversely related to the magnitude of the maternal response (Figure 3F). Highest transfer was observed after infection with the lowest maternal response and lowest transfer after vaccination and infection inducing the highest maternal response (Figure 3F). Overall transport of antibody functions across the placenta was based on the nature of the antibody function and maternal levels.

**Figure 3:**
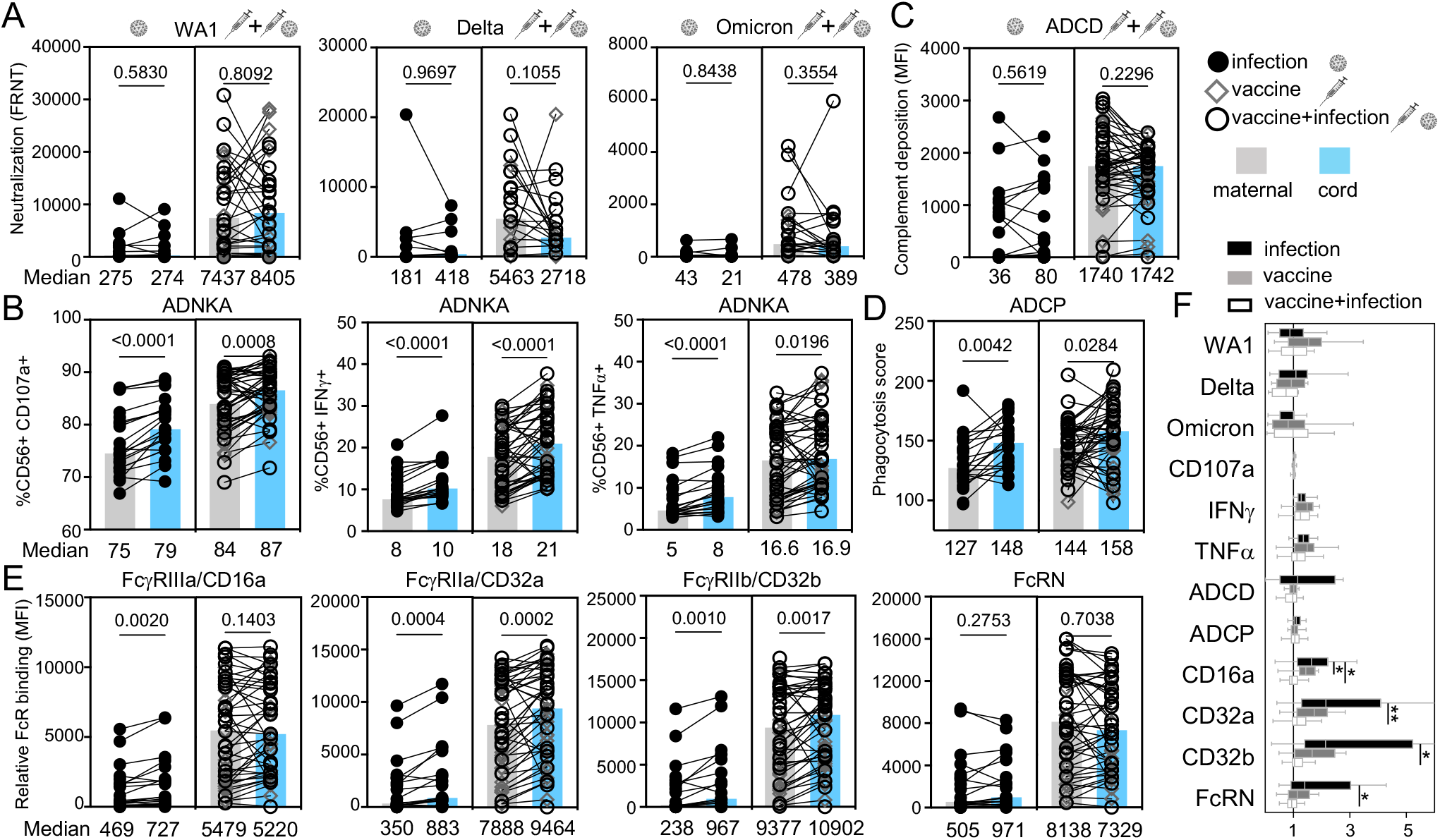
SARS-CoV-2 neutralizing and RBD Fc effector functions are differentially transferred the placenta. (**A**) Neutralization against live SARS-CoV-2 WA1, variant Delta and, (**B**) RBD ADNKA, (**C**) RBD ADCD, (**D**) RBD ADCP, and (**E**) relative binding of RBD-IgG to FcγRIIIa/CD16a, FcγRIIa/CD32a, FcγRIIb/CD32b and FcRN are compared with es of the medians for maternal (grey) and matched cord (blue) samples listed below. al significance was calculated by Wilcoxon-matched pairs test. (**F**) Antibody function ratios (the proportion of cord to maternal levels) are shown with medians (bars), rtile ranges (boxes), and ranges (whiskers).

### RBD-specific isotype and subclass

Differential isotypes and subclass drive diversity in antibody functions (36, 86). To determine which are induced by vaccination we measured RBD IgG, IgA and IgM and IgG1, IgG2, IgG3 and IgG4. In cord samples, we observed that the magnitude of RBD IgG, specifically IgG1 and not IgG2-4, was enhanced (Figure 4A), with no differences in control influenza-specific IgG (Supplemental Figure 4). RBD IgM and IgA were minimally detected (Figure 4A and Supplemental Figure 4 and 5A). Similarly, in maternal samples vaccination linked to enhanced RBD IgG, specifically IgG1 (Figure 4B). While RBD IgA and IgM were detectable and trended higher with vaccination, no differences were significant (Supplemental Figure 5B). Thus, vaccination in pregnancy enhances the magnitude of RBD IgG, specifically IgG1, in both maternal and fetal responses.

**Figure 4:**
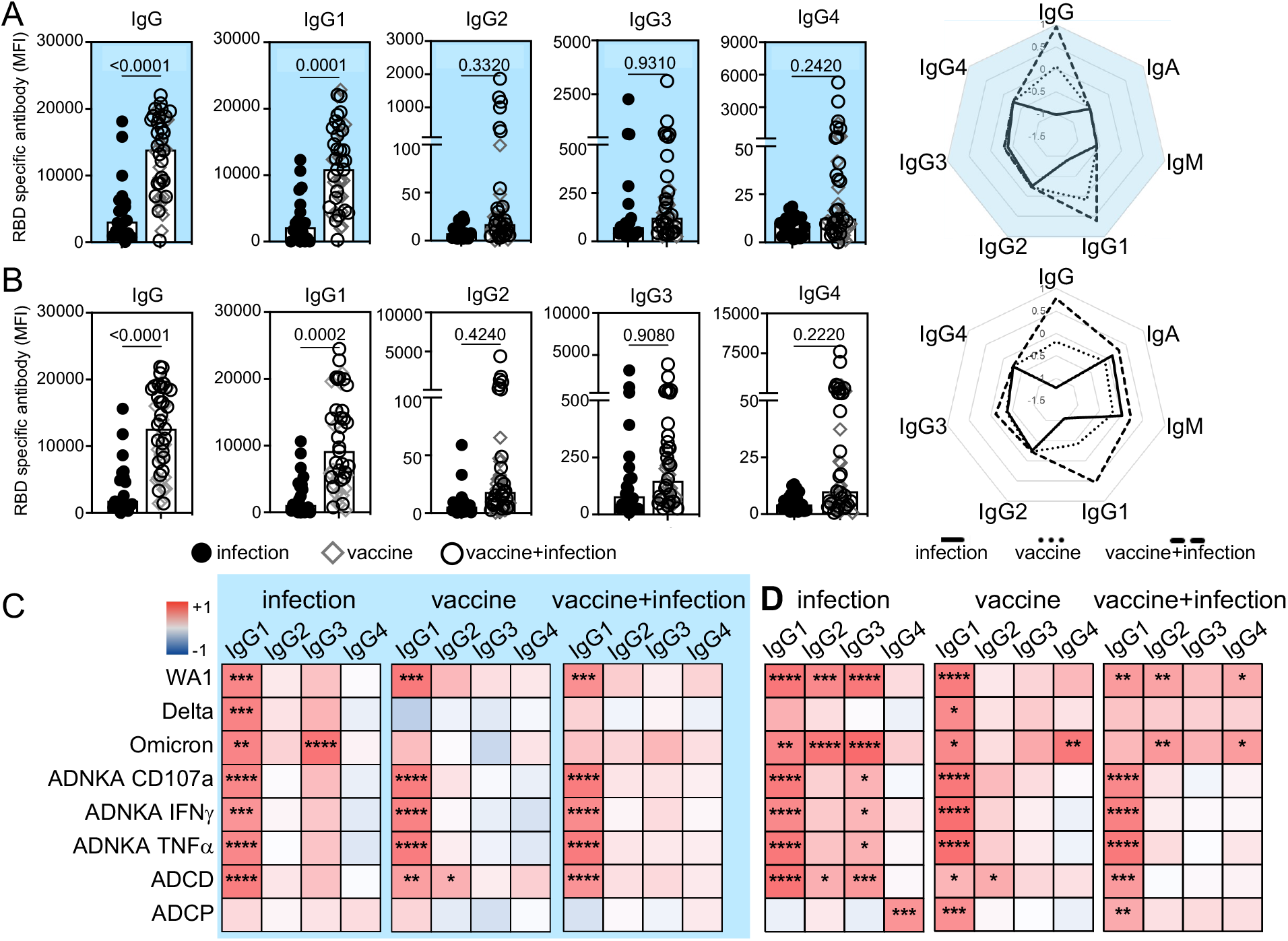
Vaccination in pregnancy enhances RBD IgG1. The magnitude of RBD-specific total subclasses in (**A**) cord and (**B**) maternal responses are shown. P-values are adjusted rnal age and body mass index using linear regression. Radar plots summarize the de of RBD-specific isotype and subclass. Each line represents the median Z-scored data clinical group (infection n=20, vaccine n=18, vaccine+infection n=27). Heatmaps of the on coefficients (r^2^) summarize the dependency of RBD-specific antibody functions on es in cord (**C**) and maternal (**D**) samples by simple linear regression. * p≤0.05; ** *** p≤0.001; **** p≤0.0001.

### IgG subclasses and antibody functions

With the increased magnitude of RBD IgG1 after vaccination, we expected this subclass to drive antibody functions targeting SARS-CoV-2. Using linear regression to assess the dependency of each neutralizing and Fc effector function on RBD-specific subclasses we found that this was the general case for maternal and cord responses across immune exposures (Figure 4C, D). A greater breadth of subclasses (RBD IgG1, IgG2, IgG3 and IgG4) was linked to antibody functions in maternal blood, which focused on IgG1 with vaccination (Figure 4D).

### RBD-specific IgG glycosylation

Differential post-translational IgG glycosylation modulates Fc effector functions in SARS-CoV-2 infection and vaccination (26, 45-48, 50) and potentially placental transfer (31, 66, 67, 87). A core bi-antennary structure of mannose and N-acetylglucosamine on a conserved N297 of the Fc domain is modified with the addition and subtraction of galactose (G), sialic acid (S), fucose (F) and a bisecting N-acetylglucosamine GlcNAc (B) (Figure 5, Supplemental Figure 6) (52) into diverse forms. We quantitated the relative abundance of the individual glycoforms (Figure 5 and Supplemental Figure 6) and found differences between RBD and non-antigen specific IgG in both maternal and cord blood (Figure 5A, B). However, vaccination increased fucosylated (F) and decreased di-sialylated (S) structures on cord RBD IgG (Figure 5C and Supplemental Figure 7A). These differences were not observed on non-antigen specific (Supplemental Figure 7B) or on maternal IgG (Figure 5D and Supplemental Figure 7C-D). Thus, RBD-specific IgG glycosylation highlights differences in maternal-cord blood with vaccination compared to infection in pregnancy.

**Figure 5:**
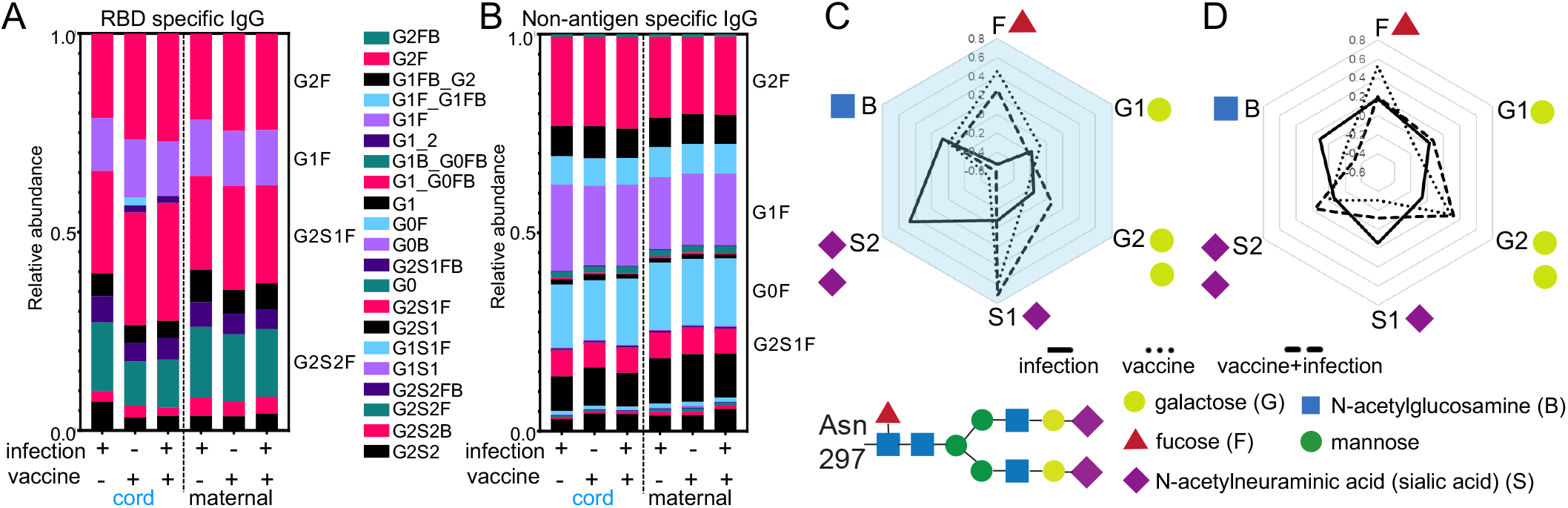
Vaccination in pregnancy changes glycosylation of infant cord and not maternal RBD-IgG. The relative abundance of (**A**) RBD and (**B**) non-antigen specific IgG individual ms are depicted by the medians in each clinical group. Radar plots summarize (**C**) cord maternal glycoforms on RBD relative to non-antigen specific IgG for each sample with wing the medians for each clinical group.

### IgG glycosylation and antibody functional potency

Antibody glycosylation impacts functional potency, the level of function relative amount of antibodies (88, 89). Of all antibody functions measured, potency of viral neutralization against WA1 and ADCD were significantly enhanced in the vaccinated compared to only infected (Figure 6A, B). No differences were observed for ADNKA and ADCP (Figure 6A, B). Thus, vaccination in pregnancy enhances the potency of a subset of maternal and fetal antibody functions.

**Figure 6:**
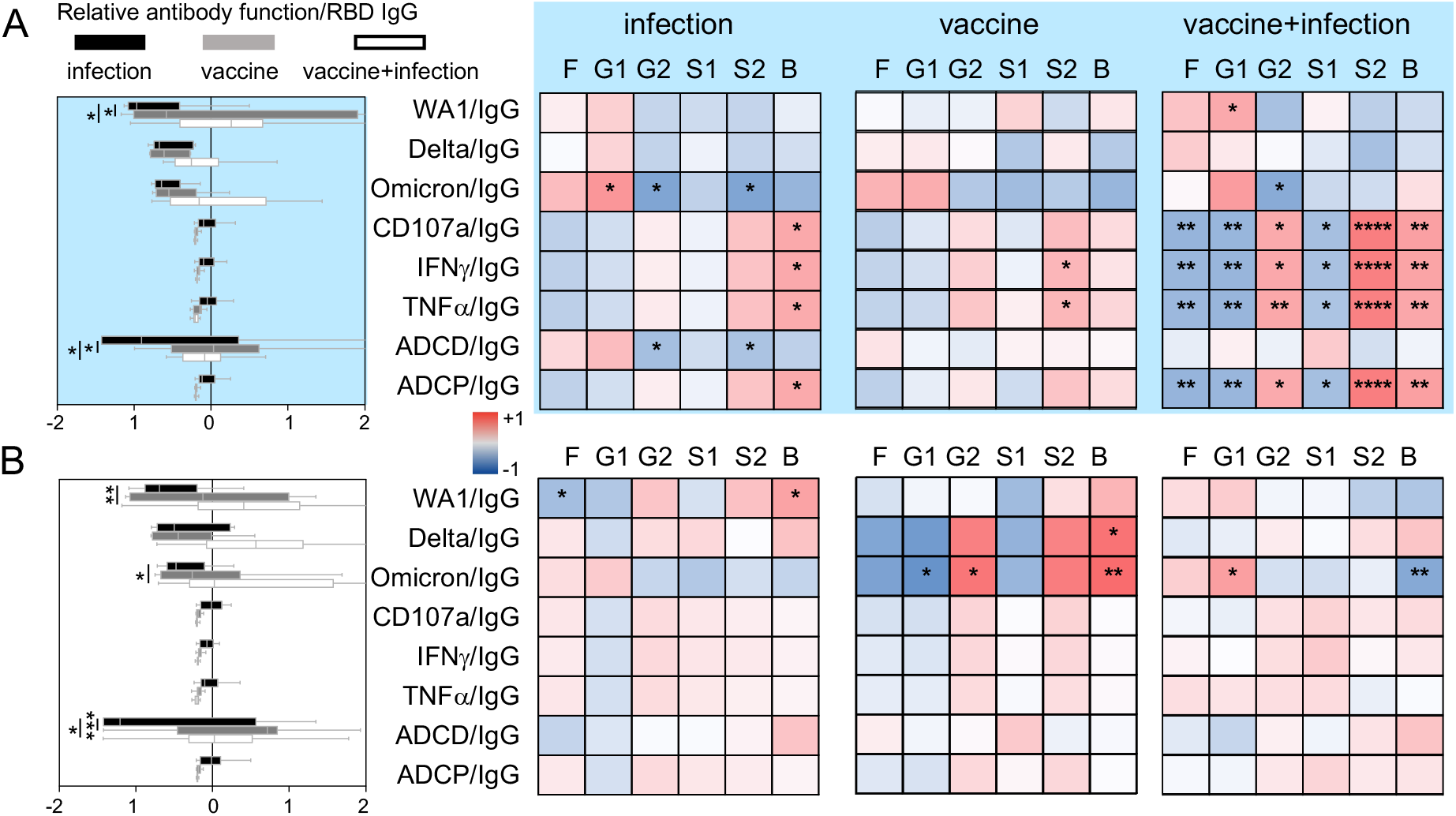
Differential RBD IgG glycosylation impacts antibody functional potency more in infant mpared to maternal responses. For every patient sample, potency was calculated for tibody function. The medians (bars), interquartile ranges (boxes), and ranges (whiskers) -scored data for (**A**, left) cord and (**B**, left) maternal samples in each clinical group are Heatmap of the regression coefficients (r^2^) summarizes the dependency of RBD-specific functions on RBD-specific IgG glycans in (**A**, right) cord and (**B**, right) maternal samples le linear regression. Fucosylated (F), monogalactosylated (G1), digalactosylated (G2), lylated (S1), disialylated (S2) and bisecting n-acetyl-glucosamine (B) glycoforms are p≤0.05; ** p≤0.01; *** p≤0.001; **** p≤0.0001.

To examine how differential RBD IgG glycans influence antibody functional potency, we used linear regression. Antibody functional potency in cord compared to maternal samples was generally more dependent on glycans (Figure 6A, B). The most prominent difference was observed in the combination vaccine and infection group, highlighting the negative effect of fucose and positive of di-sialic acid, the only RBD IgG glycans that changed significantly with vaccination compared to infection (Supplemental Figure 7A).

### Vaccine mediated polyfunctional antibody coordination

Polyclonal responses offer protection against subsequent viral challenge through a integration of multiple antibody functions (42, 55). We found partial overlap in the coordination between neutralizing and Fc effector functions in maternal and cord responses (Figure 7A). Correlations between viral neutralization and RBD ADNKA and Fcγ receptor binding were relatively preserved across the maternal-fetal dyad irrespective of immune exposure. Links to RBD ADCP were observed more in maternal samples after vaccination. In contrast, links to RBD ADCD were observed in both maternal and cord samples after infection. Finally, more differences between maternal and cord samples were apparent after vaccination as compared to natural infection, suggesting diverging responses in the context of specific immune exposures.

**Figure 7:**
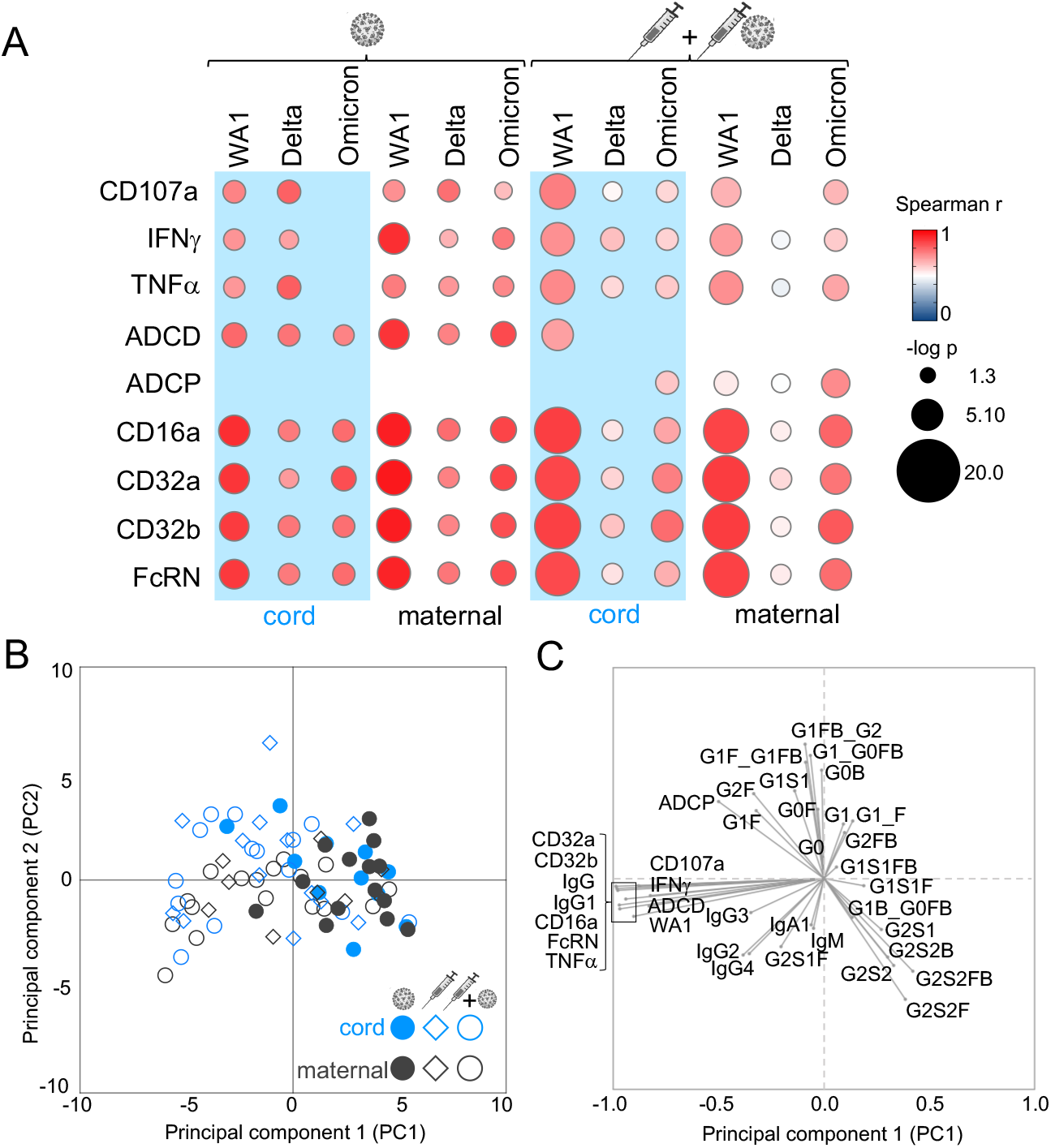
Antibody functions highlight the effect of differential immune exposure in pregnancy cosylation marks diverging maternal and infant cord responses. Bubble plots (**A**) show lation between neutralizing activities against SARS-CoV-2 WA1, Delta and Omicron and ecific Fc effector functions of antibody-dependent natural killer cell activation (CD107a, Fα), antibody-dependent complement deposition (ADCD), antibody-dependent cellular tosis (ADCP) and relative binding to FcγR (FcγRIIIa/CD16a, FcγRIIa/CD32a, CD32b, FcRN). The Spearman’s rank correlation coefficient is shown by color and nce (-log p) by size with those p<0.05 depicted. Principle-component analysis (PCA) SARS-CoV-2 antibody functions and features show separations between infection and clinical groups and maternal and cord responses. Each symbol in the (**B**) score plot ts a single maternal or cord sample. Each antibody feature is represented in the (**C**) plot, where its location reflects the distribution of the individual samples in the (**B**) score plot

To globally assess responses by maternal-fetal origin and immune exposure, we performed principal component analysis using all 38 measured antibody features with the capacity to modify SARS-CoV-2 in subsequent challenge. We found partial overlap between vaccine and infection groups and within the maternal-fetal dyad (Figure 7B, C). Explaining the most amount of variance in the data through PC1 were the top antibody features that distinguished the vaccine from infection group-RBD Fcγ receptor binding and IgG levels-affirming our univariate analyses (Figure 1, 2, 4). Explaining the second most amount of variance through PC2 was RBD-specific antibody glycosylation, driving differences between maternal and cord samples. Thus, vaccination in pregnancy enriches multiple antibody neutralizing and Fc functions with glycosylation differentiating fetal from maternal responses.

## DISCUSSION

### Neutralizing and antibody Fc effector functions transfer to the fetus

Observational studies show that two doses of monovalent mRNA vaccines in pregnancy protect infants up to 6 months after birth from complications and morbidity associated with COVID-19 but what mediates protection is less clear (15, 19). The data here shows that in cord blood vaccination compared to infection enhances overall magnitude, polyfunctional breadth (Figure 1 and 2) and antibody potency (Figure 6) against SARS-CoV-2 through RBD IgG (Figure 4 and Supplemental Figure 5), specifically IgG1 (Figure 4) and differential antibody glycosylation (Figure 5-7) that drives coordinated (Figure 7A). These include neutralization of SARS-CoV-2 WA1 (Figure 1A, F) and binding of RBD-specific antibodies to Fcγ receptors (Figure 1E, F) involved in the induction of natural killer cell activation and concomitant cellular cytotoxicity (Figure 1B, F). Complementary to Fc receptor binding, C1q engagement of the antibody Fc domain to induce C3 mediated complement activation is enhanced, offering protection via a separate mechanism from neutralizing and other Fcγ effector functions (Figure 1C, 7A). Not all functions are enriched in vaccination compared to infection as in the case of cord antibody-dependent cellular phagocytosis (Figure 1D, 7). However, Fcγ receptor mediated functions compared to neutralizing and complement activities are preferentially transported to the fetus regardless of whether the immune exposure is infection or vaccination (Figure 3). Thus, a subset of vaccine-induced maternal antibody neutralizing and Fc effector functions is transferred to the fetus.

Accumulating data suggest that immune responses beyond direct neutralization likely shape protection in subsequent viral challenge (20-25, 55). In animal models testing monoclonal antibodies and vaccination, the absence of Fc domain-Fcγ receptor engagement leads to increased viral load and pathology (25, 38-42). As such, preserved Fc effector functions (25, 26, 43) likely do not prevent acquisition of infection but rather limit disease in the absence of high neutralization as is observed with the monovalent WA1-based mRNA vaccines against Delta and Omicron (90, 91).

Fc effector functions may be particularly relevant for the fetus. Similar to HIV, CMV and malaria and some vaccines (67-69, 92), the data from this study show that Fcγ receptor mediated functions are transferred along with IgG across the placenta (Figure 1, 3). Higher natural killer cell activation and cellular phagocytosis in cord compared to paired maternal samples (Figure 3B, D) irrespective of immune exposure suggest preferential transport of Fcγ functions that contrast equivalent levels of neutralizing (Figure 3A) and complement (Figure 3C) activities. Longitudinal studies show that Fc functions persist longer than neutralizing activities (23, 29, 78). Thus, Fc functions have the localization and durability to offer protection across the gestational and neonatal periods when the immune system is immature.

### Vaccine-induced maternal antibody functions transfer to the fetus

The data from this study show that vaccination as compared to a spectrum of SARS-CoV-2 infection from asymptomatic to severe COVID-19 (Supplemental Figure 1) in pregnancy enhances maternal antibody neutralizing and Fc effector functions. Mirroring paired cord responses, neutralization against WA1, Fcγ receptor binding mediated natural killer cell activation and C1q-C3 driven complement activation are elevated in maternal responses to vaccination compared to infection alone (Figure 2, Figure 3A-C). This occurs even in the context of co-morbidities including pre-gestational diabetes and chronic hypertension (Supplemental Table 1). Thus, like the fetus, maternal vaccination likely also benefits the pregnant individual through enhanced neutralizing and Fc effector functions.

Antibodies and their associated functions traffic across the placenta. The primary transporter of IgG from the pregnant individual to the fetus is the neonatal FcRN (80, 81), though FcγRIIIa/CD16a, FcγRIIa/CD32a, FcγRIIb/CD32b and other Fcγ receptors expressed in the placenta could be additional factors (31, 67). Consistent with its high affinity nature, relative binding to FcRN is equivalent in cord and maternal responses irrespective of immune exposure (Figure 3E). Yet with antibodies binding to the low affinity Fcγ receptors there is preferential fetal localization when the maternal magnitude is low (Figure 3E, F). Upon vaccination, the difference in FcγRIIIa/CD16a but neither FcγRIIa/CD32a nor FcγRIIb/CD32b disappears (Figure 3E), suggesting that when maternal antibodies are elevated, FcγRIIIa/CD16a binding is saturated. This effect is more apparent when maternal antibodies are highest after the combination of infection and vaccination (Figure 3F). Altered placental IgG transfer has been described in chronic infections such as HIV (67) and SARS-CoV-2 infection in pregnancy (31). Thus it is plausible that vaccination could similarly impact FcR levels and the transfer of distinctly glycosylated IgG. However, the data here shows that the large enhancement of the initial maternal response with vaccination compared to infection overshadows the smaller effects of differential transfer.

### Differential glycosylation contributes to antibody functional potency in the fetus

While vaccine-induced RBD IgG1 drives functions across the maternal-fetal dyad (Figure 4C, D), post-translational antibody glycosylation appears to contribute more to potency in the fetus than the pregnant individual (Figure 5, 6, 7B, 7C, Supplemental Figure 7 and 8). All polyclonal IgG have N-linked glycans on the Fc domain that drive Fcγ receptors binding and effector functions (*38, 39, 54, 88)* and 20% of the population have additional modifications on the Fab domain (93, 94) that have the potential to impact stability, half-life and avidity to antigens (59, 61, 95). Subtle differences between maternal and cord IgG glycosylation have been reported (87, 96). In COVID-19 disease and upon mRNA vaccination, changes in fucose and sialic acid alter antibody-dependent natural killer cell activity that leads to cellular cytotoxicity and are associated with different clinical outcomes (45-48, 97, 98). Our data show that these same glycans are altered on cord and not maternal RBD IgG with vaccination (Figure 5 and Supplemental Figure 7), impacting RBD ADNKA potency in the newborn (Figure 6). Recent data in a mouse model of Listeria infection show that vaccination during as compared to before pregnancy induces Fab domain sialylation that enhances protection for pups (61). Whether pregnancy-specific changes in antibody glycosylation could similarly improve infant protection against COVID-19 is not known. Beyond COVID-19, different vaccine adjuvants and platforms can modulate antibody glycosylation as well as subclass that could impact Fc effector functions (99, 100). Thus, what is learned in COVID-19 has implications for the design of vaccines across infections that disproportionately impact the maternal-fetal dyad (62, 68, 101, 102).

The mechanism by which IgG is differentially glycosylated between the maternal and fetal sides of the placenta are not known. One explanation could be differential transfer (31, 66, 103, 104). Another is that B cell extrinsic glycosidases and glycosyltransferases could modify antibodies as they traffic from the pregnant individual to the fetus (105, 106). Because immunity directly passed to the fetus is primarily IgG, this single isotype is the main source of protection and differential glycosylation a significant determinant of diversity. Pregnant individuals have a greater breadth of responses to subsequent viral challenge with isotypes and cellular immunity (22, 107, 108) that could leave IgG post-translational modifications less critical.

### Maternal vaccination enriches antibody functions with and without SARS-CoV-2 infection

In the general population, the combination of vaccine and infection boosts antibody responses compared to either immune exposure alone (56, 85, 109, 110). This similarly occurs in pregnancy where levels of antibody functions, Fc receptor binding and subclasses with the combination is statistically significantly higher compared to infection alone and non-significantly higher compared to vaccine alone (Figure 1, 2). Thus, maternal immunization likely benefits the pregnant individual and the fetus regardless of the presence of infection-derived immunity.

The preponderance of data show that immunity from vaccination compared to SARS-CoV-2 infection provides more effective protection against disease across different populations including the immunocompromised and the elderly. COVID-19 vaccines may be an annual recommendation for the general population but how these tools can be best leveraged with respect to trimester of immunization (33), vaccine platforms (27), and adjuvants (100) that alter antibody glycosylation and potentially functions remains to be seen. Moreover, maternal co-morbid conditions that affect the immune substrate such as gestational diabetes could impact how antibody functions are transferred to the fetus (111). Finally, newborn immunity can be further supported by maternal IgA, IgG and IgM in the colostrum (112). Studies powered to address how these factors influences antibody functions will illuminate ways to synergize pre- and post- natal immunization strategies to optimally protect the maternal-fetal dyad (113).

With no evidence of serious mRNA vaccine related adverse effects in pregnancy (114-116) and the potential benefits of maternal antibodies in preventing disease for the maternal-fetal dyad, pregnancy is argued to be a condition that should be eligible for additional doses (117). That infants under 6 months of age are the only ones with no available COVID-19 vaccines further supports this argument. However, the transfer of maternal antibodies to the fetus has many potential consequences yet to be unraveled. Some data suggest that maternal antibodies blunt infant responses to vaccines given after birth via mechanisms still debated (118-121) while others show a priming effect on infant cellular and humoral responses (122). Beyond the infant, data from animal models suggest that maternal antibodies can shape the B cell repertoire of the offspring long after the maternal antibodies themselves become undetectable (123), suggestive of a “vaccinal“ effect (124). Thus, diverging maternal and cord antibody functions from SARS-CoV-2 infection and vaccination in pregnancy could have implications for the development of immune responses to subsequent coronavirus challenges reaching beyond infancy and into adulthood.

## MATERIALS AND METHODS

### Study design and participant recruitment

We approached individuals at Parkland Health, the Dallas County public hospital in Texas. Eligible participants were pregnant, ≥18 years, able to provide informed consent, and received at least one dose of mRNA vaccine and or were infected with SARS-CoV-2 in pregnancy. Individuals with infection or vaccination outside of pregnancy were excluded. Clinical COVID-19, pregnancy and obstetric data were determined through electronic health record review. Disease severity was classified per NIH COVID-19 guidelines (125).

### Study Approval

This study was conducted in accordance with the UTSW and Parkland Health IRBs (STU2020-0375, STU2020-0214) and approved by the Oregon Health & Science University IRB (PROTO202000015). Written informed consent was received from all study individuals prior to participation.

### Sample collection

Paired maternal and infant cord blood were collected in deliveries (February 17, 2021 - May 27, 2022) by venipuncture or from umbilical vein in ACD and SST tubes. Plasma and serum were isolated by centrifugation at 1400RPM and 3400RPM respectively for 10minutes at RT, aliquoted into cryogenic vials, stored at -80°C, and heat-inactivated at 55°C for 10minutes prior to use in assays.

### Cell lines

Vero E6 cells (ATCC VERO C1008) were grown at 37°C, 5% CO2 in Dulbecco’s Modified Eagle Medium supplemented with 10% fetal bovine serum, 1% penicillin/streptomycin, 1% non-essential amino acids. THP-1 cells (ATCC TIB-202) were grown at 37°C, 5% CO2 in RPMI-1640 supplemented with 10% fetal bovine serum, 2mM L-glutamine, 10mM HEPES, and 0.05mM β-mercaptoethanol. CD16.NK-92 (ATCC PTA-6967) were grown at 37°C, 5% CO2 in MEM-α supplemented with 12.5% FBS, 12.5% horse serum, 1.5g/L sodium bicarbonate, 0.02mM folic acid, 0.2mM inositol, 0.1mM 2-β-mercaptoethanol, 100U/mL IL-2.

### Viruses

SARS-CoV-2 clinical isolates were passaged once before use: USA-WA1/2020 [original strain] (BEI Resources NR-52281); hCoV-19/USA/PHC658/2021 [B.1.617.2] (BEI Resources NR-55611), and hCoV-19/USA/CO-CDPHE-2102544747/2021 [B.1.1.529 - BA.2] (BEI Resources NR-56520). Isolates were propagated in Vero E6 cells for 24-72hours until cultures displayed at least 20% cytopathic effect (CPE) (97).

### Focus Reduction Neutralization Test (FRNT)

Focus forming assays were performed as described (26, 126). Sub-confluent Vero E6 cells were incubated for 1hour with 30µL of diluted sera (5x4-fold starting at 1:20) which was pre-incubated for 1hour with 100 infectious viral particles per well. Samples were tested in duplicate. Wells were covered with 150µL of overlay media containing 1% methylcellulose and incubated for 24hours. Plates were fixed by soaking in 4% formaldehyde in PBS for 1hour at RT. After permeabilization with 0.1% BSA, 0.1% saponin in PBS, plates were incubated overnight at 4°C with primary antibody (1:5,000 anti-SARS-CoV-2 alpaca serum) (Capralogics Inc) (126). Plates were then washed and incubated for 2hours at RT with secondary antibody (1:20,000 anti-alpaca-HRP) (NB7242 Novus) and developed with TrueBlue (SeraCare) for 30minutes. Foci were imaged with a CTL Immunospot Analyzer, enumerated using the viridot package (127) and %neutralization calculated relative to the average of virus-only wells for each plate. FRNT50 values were determined by fitting %neutralization to a 3-parameter logistic model as described previously (126). The limit of detection (LOD) was defined by the lowest dilution tested, values below the LOD were set to LOD – 1. Duplicate FRNT50 values were first calculated separately to confirm values were within 4-fold. When true, a final FRNT50 was calculated by fitting to combined replicates.

### Antigen-specific antibody isotype and subclass

Quantification of antigen-specific IgG and subclasses, IgM, and IgA1 was performed as described (26, 128). Carboxylated microspheres (Luminex) were coupled with recombinant SARS-CoV-2 RBD (129) (BEI Resources NR-52309) by covalent NHS-ester linkages via EDC (1-Ethyl-3-[3-dimethylaminopropyl] carbodiimide hydrochloride) and Sulfo-NHS (N-hydroxysulfosuccinimide) (ThermoScientific) per manufacturer instructions. A mixture of influenza antigens from strain H1N1 (NR-20083 and NR-51702, BEI Resources), H5N1 (NR-12148, BEI Resources), H3N2, B Yamagata lineage, and B Victoria lineage (NR-51702, BEI Resources) was used as a control. Antigen-coupled microspheres (1000/well) were incubated with serially diluted samples (IgG at 1:100, 1:1000, 1:10,000; IgM at 1:100, 1:300, 1:900; IgA1 at 1:30, 1:90, 1:270) in replicates in Bioplex plates (Bio-Rad) at 4°C for 16hours. After washing away the unbound antibodies, bead bound antigen-specific antibodies were detected by using PE-coupled detection antibody (anti-IgG, IgA1, IgM, IgG1, IgG2, IgG3 and IgG4 from Southern Biotech) (1µg/mL). After 2hours of incubation at RT, the beads were washed with PBS 0.05% Tween20 and PE signal measured on a MAGPIX (Luminex). The background signal (PBS) was subtracted. Experiments were conducted two independent times. Representative data from one dilution was chosen by the highest signal-to-noise ratio.

### Antigen-specific antibody Fc receptor binding

Relative Fc receptor binding was assessed as described (26, 130). Luminex carboxylated microspheres were coupled with antigens as described for antigen subclass and isotype above. Antigen-coupled microspheres (1000/well) were incubated with serially diluted samples (1:100, 1:1000, 1:10,000) in replicates in Bioplex plates (Bio-Rad) at 4°C for 16hours. Recombinant Fc receptors (FcγRIIIa/CD16a, FcγRIIa/CD32a H167, FcγRIIb/CD32b, Neonatal Fc receptor/FcRN) (R&D Systems) were labeled with PE (Abcam) per manufacturer’s instructions, added (1µg/mL) to bead bound antigen-specific immune complexes. After 2hours of incubation at RT, the beads were washed and antigen-specific antibody bound Fc receptors were measured on MAGPIX (Luminex). The background signal (PBS) was subtracted. Experiments were conducted two independent times. Representative data from one dilution was chosen by the highest signal-to-noise ratio.

### Antigen-specific antibody-dependent cellular phagocytosis (ADCP)

The THP-1 (TIB-202, ATCC) ADCP with antigen-coated beads was conducted as described (26). SARS-CoV-2 RBD (BEI Resources NR-52309) was biotinylated with Sulfo-NHS-LC Biotin (Thermo Fisher), then incubated with 1μm fluorescent neutravidin beads (Invitrogen) at 4°C for 16hours. Excess antigen was washed away and RBD-coupled neutravidin beads were resuspended in PBS-0.1% bovine serum albumin (BSA). RBD-coupled beads were incubated with serially diluted samples (1:100, 1:500, 1:2500) in duplicate for 2hours at 37°C. THP1 cells (1×10^5^ per well) were then added. Serum opsonized RBD-coupled beads and THP1 cells were incubated at 37°C for 16hours, washed and fixed with 4% PFA. Bead uptake was measured on a BD LSRFortessa and analyzed by FlowJo10. Phagocytic scores were calculated as the integrated median fluorescence intensity (MFI) (%bead-positive frequency×MFI/10,000) (131). The background signal (PBS) was subtracted. Experiments were conducted two independent times. Representative data from one dilution was chosen by the highest signal-to-noise ratio.

### Antibody-dependent complement deposition (ADCD)

The ADCD assay was performed as described (26, 132). Luminex carboxylated microspheres were coupled with antigens as described for antigen subclass and isotype above. Antigen-coated microspheres (2500/well) were incubated with serially diluted heat inactivated samples (1:10, 1:50, 1:250) at 37°C for 2hours. Guinea pig complement (Cedarlane) freshly diluted 1:60 in PBS was added for 20minutes at 37°C. After washing off excess with PBS 15mM EDTA, anti-C3 PE-conjugated goat polyclonal IgG (MP Biomedicals) (1µg/mL) was added. The beads were then washed and C3 deposition quantified on a MAGPIX (Luminex). The background signal (PBS) was subtracted. Experiments were conducted two independent times. Representative data from one dilution was chosen by the highest signal-to-noise ratio.

### Antibody-dependent NK cell activation (ADNKA)

ADNKA was performed as described (26, 133). ELISA plates were coated with recombinant RBD (300 ng/well) (BEI Resources NR-52309). Wells were washed, blocked, and incubated with serially diluted samples (1:10, 1:100, 1:1000) in duplicate for 2hours at 37°C prior to adding CD16a.NK-92 cells (PTA-6967, ATCC) (5 × 10^4^ cells/well) for 5hours with brefeldin A (Biolegend), Golgi Stop (BDBiosciences) and anti-CD107a (clone H4A3, BDBiosciences). Cells were stained with anti-CD56 (clone 5.1H11, BDBiosciences) and anti-CD16 (clone 3G8, BDBiosciences) and fixed with 4%PFA. Intracellular cytokine staining to detect IFNγ (clone B27, BDBiosciences) and TNFα (clone Mab11, BDBiosciences) was performed in permeabilization buffer (Biolegend). Markers were measured using a BD LSRFortessa and analyzed by FlowJo10. CD16 expression was confirmed. NK cell degranulation and activation were calculated as %CD56+CD107a+, IFNγ+ or TNFα+. Representative data from one dilution was chosen by the highest signal-to-noise ratio. Experiments were conducted two independent times.

### Non-antigen and RBD-specific IgG glycosylation

Non-antigen and RBD-specific IgG glycans were purified and relative levels were quantified as described with modifications (26, 94, 134). RBD (BEI Resources NR-52309) was biotinylated with sulfosuccinimidyl-6-[biotinamido]-6-hexanamido hexanoate (sulfo-NHS-LC-LC biotin; ThermoScientific) and coupled to streptavidin beads (New England Biolabs). Patient samples were incubated with RBD-coupled beads and excess sera washed off with PBS (Sigma). RBD-specific antibodies were eluted from beads using 100mM citric acid (pH 3.0) and neutralized with 0.5M potassium phosphate (pH 9.0). Non-antigen specific IgG and RBD-specific IgG were isolated from serum and eluted RBD-specific antibodies respectively by protein G beads (Millipore). Purified IgG was denatured and treated with PNGase enzyme (New England Biolabs) for 12hours at 37°C to release glycans.

To isolate bulk IgG glycans, proteins were removed by precipitation using ice cold 100% ethanol at -20°C for 10minutes. To isolated RBD-specific IgG glycans, Agencourt CleanSEQ beads (Beckman Coulter) were used to bind glycans in 87.5% acetonitrile (Fisher Scientific). The supernatant was removed, glycans were eluted from beads with HPLC grade water (FisherScientific) and dried by centrifugal force and vacuum (CentriVap). Glycans were fluorescently labeled with a 1.5:1 ratio of 50mM APTS (8-aminoinopyrene-1,3,6-trisulfonic acid, ThermoFisher) in 1.2M citric acid and 1M sodium cyanoborohydride in tetrahydrofuran (FisherScientific) at 55°C for 3hours. Labeled glycans were dissolved in HPLC grade water (FisherScientific) and excess unbound APTS removed using Agencourt CleanSEQ beads (non-antigen specific glycans) and Bio-Gel P-2 (Bio-rad) size exclusion resin (RBD-specific glycans). Glycan samples were run with a LIZ 600 DNA ladder in Hi-Di formamide (ThermoFisher) on an ABI 3500xL DNA sequencer and analyzed with GlycanAssure Data Acquisition Software v.1.0. Each glycoform was identified by standard libraries (GKSP-520, Agilent). The relative abundance of each glycan was determined as the proportion of each individual peak with respect to all captured.

### Statistical analyses

Statistical analyses were performed using R4.1.2, Stata17 and GraphPad 9.0. Data are summarized using median (Q1-Q3), mean ± standard deviation, percent (%). Data were evaluated for normality and independence of the residuals, heteroscedasticity and linear relationships between dependent and independent variables and log transformed as needed to meet these assumptions. Linear regression models were used to adjust for the effect of maternal age and BMI (Figure 1-2, 4, Supplemental Figure 4-5, 7) and determine the effects of fetal sex and disease severity. Wilcoxon-matched pair signed rank tests were used to compare between maternal-cord pairs (Figure 3). For antibody function radar plots (Figure 1F, 2F, 4), Z-scored data for each feature were calculated and the median values for each group plotted. For the antibody glycan radar plots (Figure 5, C and D), Z scores of individual RBD-specific relative to non-antigen specific IgG glycoforms were calculated and the medians for each group plotted. Simple linear regression was used to examine the relationships between IgG subclass as the independent and antibody functions as the dependent variables (Figure 4, C and D) and IgG glycoforms as the independent and antibody functional potencies as the dependent variables (Figure 6). Spearman rank correlations were used to examine bivariate associations between antibody functions (Figure 7A). Principal component analysis (135) was used to reduce variable dimensions (Figure 7B-C, Supplemental Figure 8). For clinical data in Tables, analysis of variance was used for age as it was normally distributed, Kruskal-Wallis test was used for all other continuous variables and Chi-square and Fisher’s exact tests for categorical variables. All p-values are two-sided, and <0.05 considered significant. In Figures, asterisks denote statistical significance (∗ p≤0.05; ∗∗ p≤0.01; ∗∗∗ p≤0.001; ∗∗∗ p≤0.0001).

## Author contributions

EA obtained patient samples, collected clinical data, and prepared the manuscript. PL designed and conducted experiments, analyzed data, and prepared the manuscript. YJK assisted in conducting experiments, analyzing data, and preparing the manuscript. AM and JP analyzed the data. TAB, MGT, and SKM conducted the neutralization assays and analyzed data. FGT supervised neutralization assays, analyzed data, and provided critical revisions to the manuscript. LLL conceived, designed, coordinated, and supervised the work, analyzed the data, and wrote the manuscript.

## Acknowledgements

We thank the patients, Brenda Espino, and the UTSW COVID-19 Biorepository (Dwight Towler, David Greenberg, Benjamin Greenberg, and Nancy Monson) for samples. We thank Catherine Spong and Trish Perl for their support. Gabrielle Lessen and Joshua Miles provided graphical assistance. Funding is provided by UTSW Internal Medicine Pilot Project Award (E.H.A. and L.L.L.), Disease Oriented Clinical Scholars Award (L.L.L.), Burroughs-Wellcome Fund UTSW Training Resident Doctors as Innovators in Science (Y.J.K.), NIH T32HL083808 (T.A.B.), NIH R011R01AI141549-01A1 (F.G.T.), OHSU Innovative IDEA grant 1018784 (F.G.T.), UT-FOCUS through American Heart Association (E.H.A.), Doris Duke Charitable Foundation (E.H.A.), the Harry S. Moss Heart Trust (E.H.A.), and Parkland Community Health Plan, a component unit of Dallas County Hospital District d/b/a Parkland Health.

---

**Supplemental Table 1.** Additional clinical characteristics of study patients

| **Characteristic** | **Infection**<br>**n = 22** | **Vaccine**<br>**n = 19** | **Vaccine**<br>**and**<br>**infection**<br>**n = 28** | **P value** |
| --- | --- | --- | --- | --- |
| Nulliparous | 9 (41) | 5 (26) | 3 (11) | 0.0481 |
| Pregestational diabetes | 3 (14) | 2 (11) | 8 (29) | 0.266 |
| Chronic hypertension | 3 (14) | 5 (26) | 4 (14) | 0.562 |
| Preeclampsia with severe features | 3 (14) | 2 (11) | 7 (25) | 0.444 |
| Chorioamnionitis | 2 (9) | 0 (0) | 1 (4) | 0.481 |
| Prelabor rupture of membranes | 2 (9) | 0 (0) | 5 (18) | 0.139 |
| Induction of labor | 13 (59) | 8 (42) | 12 (43) | 0.439 |
| Cesarean delivery | 7 (32) | 11 (58) | 13 (46) | 0.241 |
| EGA <37 weeks at delivery | 7 (32) | 4 (21) | 6 (21) | 0.684 |
| Infant birth weight <10<sup>th</sup> percentile | 2 (9) | 3 (16) | 5 (18) | 0.690 |
| Neonatal intensive care unit admission | 4 (18) | 0 (0) | 5 (18) | 0.130 |

Data shown as n (%), mean ± standard deviation (SD), or median (Q1-Q3) as appropriate.
BMI= body mass index, EGA= estimated gestational age, NICU= neonatal intensive care unit

**Supplemental Table 2:** Comparison of clinical characteristics of individuals receiving mRNA-1273 and NT1262b2

| **Characteristic** | **mRNA-1273**<br>**n = 4** | **BNT1262b2**<br>**n = 43** |
| --- | --- | --- |
| Age, years | 36.8 (4.7) | 33.3 (6.4) |
| Race/ethnicity |  |  |
| Hispanic | 4 (100) | 41 (95) |
| Black, non-Hispanic | 0 (0) | 1 (2) |
| White, non-Hispanic | 0 (0) | 1 (2) |
| Other | 0 (0) | 0 (0) |
| Nulliparous | 0 (0) | 8 (19) |
| BMI at first visit, kg/m<sup>2</sup> | 37.5 (34.5-39.6) | 32.1 (28.5-36.4) |
| SARS-CoV-2 in pregnancy | 3 (75) | 25 (58) |
| mRNA Vaccination in pregnancy booster | 2 (50) | 7 (16) |
| Pregestational diabetes | 2 (50) | 8 (19) |
| Chronic hypertension | 1 (25) | 8 (19) |
| Preeclampsia with severe features | 2 (50) | 7 (16) |
| Chorioamnionitis | 0 (0) | 1 (2) |
| Induction of labor | 1 (25) | 19 (44) |
| Cesarean | 3 (75) | 21 (49) |
| EGA at delivery, weeks | 29.9 (27.5-33.2) | 25.4 (16.8-29.8) |
| EGA<37 weeks delivery | 2 (50) | 8 (19) |
| Infant birth weight <10<sup>th</sup> percentile | 2 (50) | 6 (14) |
| NICU admission | 1 (25) | 4 (9) |
| Male infant sex | 1 (25) | 26 (60) |

Data shown as n (%), mean ± standard deviation (SD), or median (Q1-Q3) as appropriate.
BMI= body mass index, EGA= estimated gestational age, NICU= neonatal intensive care unit

**Supplemental Table 3.** Characteristics of study cohort compared with deliveries at Parkland Health, 021

| **Characteristic** | **Study patients**<br>**n = 69** | **Deliveries 2021**<br>**n = 11,170** |
| --- | --- | --- |
| Age, years | 31.9 ± 6.8 | 27.5 ± 6.4 |
| Race/ethnicity |  |  |
| Hispanic | 65 (94) | 8643 (77) |
| Black, non-Hispanic | 3 (4) | 1799 (16) |
| White, non-Hispanic | 1 (1) | 448 (4) |
| Other |  | 280 (3) |
| Nulliparous | 17 (25) | 3431 (31) |
| BMI at first visit, kg/m<sup>2</sup> | 32 (28-37) | 29 (25-33) |
| Documented SARS-CoV-2 in pregnancy | 50 (72) | 2681 (24) |
| Any mRNA vaccination in pregnancy | 47 (68) | 2772 (25) |
| Pregestational diabetes | 13 (19) | 210 (2) |
| Chronic hypertension | 12 (17) | 847 (8) |
| Preeclampsia with severe features | 12 (17) | 1158 (10) |
| Chorioamnionitis | 3 (4) | 911 (8) |
| Prelabor rupture of membranes | 7 (10) | 2444 (22) |
| Induction of labor | 33 (48) | 3047 (27) |
| Cesarean delivery | 31 (45) | 3248 (29) |
| EGA at delivery, weeks | 38 (37-38) | 39 (38-40) |
| EGA <37 weeks at delivery | 17 (25) | 1057 (9) |
| Infant birth weight <10<sup>th</sup> percentile | 10 (14) | 1173 (11) |
| NICU admission | 9 (13) | 572 (5) |
| Male infant sex | 39 (57) | 5774 (52) |

Data shown as n (%), mean ± standard deviation (SD), or median (Q1-Q3) as appropriate.
BMI= body mass index, EGA= estimated gestational age, NICU= neonatal intensive care unit

## Notes

**Conflict-of-interest statement:** The authors have declared that no conflict of interest exists.

### Competing Interest Statement

The authors have declared no competing interest.

