## supplemental figures for "Diverging maternal and infant cord antibody functions from SARS-CoV-2 infection and vaccination in pregnancy"

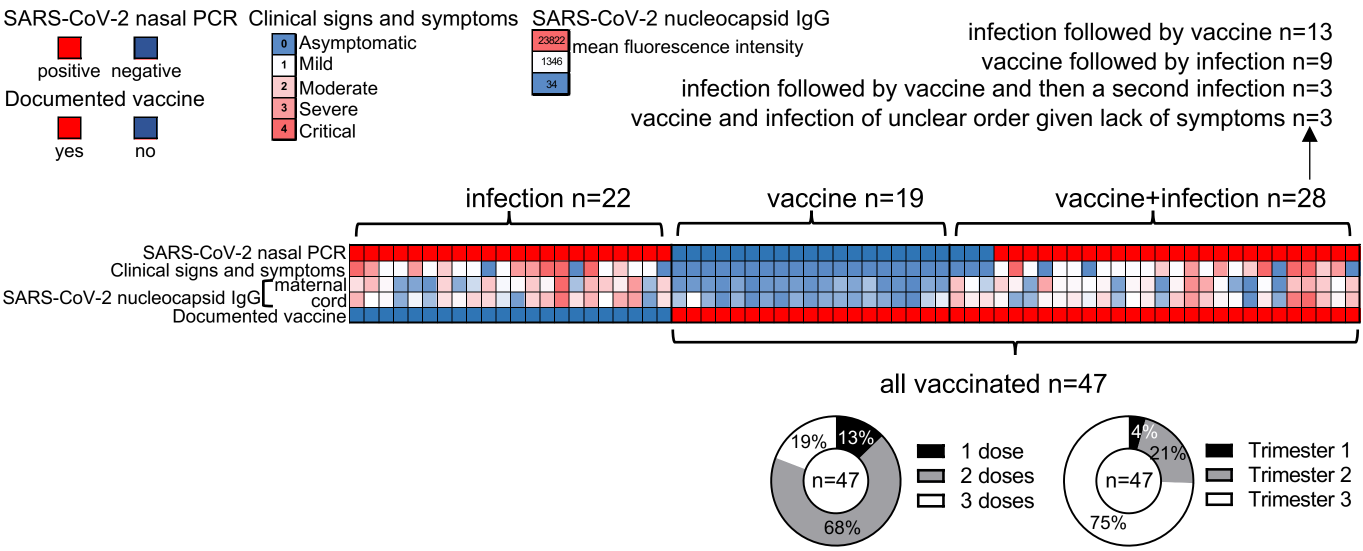


Supplemental Figure 1: Laboratory and clinical data were used to define infection and vaccine groups. Heatmap depicts results from SARS-CoV-2 nasal PCR, clinical signs and symptoms with respect to COVID-19 from asymptomatic to critical disease ascertained by chart review, SARS-CoV-2 nucleocapsid IgG in maternal and cord samples and documentation of BNT162b2 or mRNA-1273 vaccination during pregnancy in the medical record. Disease severity in pregnancy was classified as asymptomatic (no symptoms), mild (upper respiratory or mild febrile illness without lower respiratory symptoms), moderate (lower respiratory symptoms without oxygen requirement, with SpO2 ≥94% on room air), severe (oxygen requirement, or SpO2 <94% on room air), or critical (respiratory failure, mechanical ventilation, or ECMO). Pie charts depict the percentage of individuals received vaccine doses (left chart) and the percentage of individuals received last dose vaccine during three trimesters before delivery (right chart).

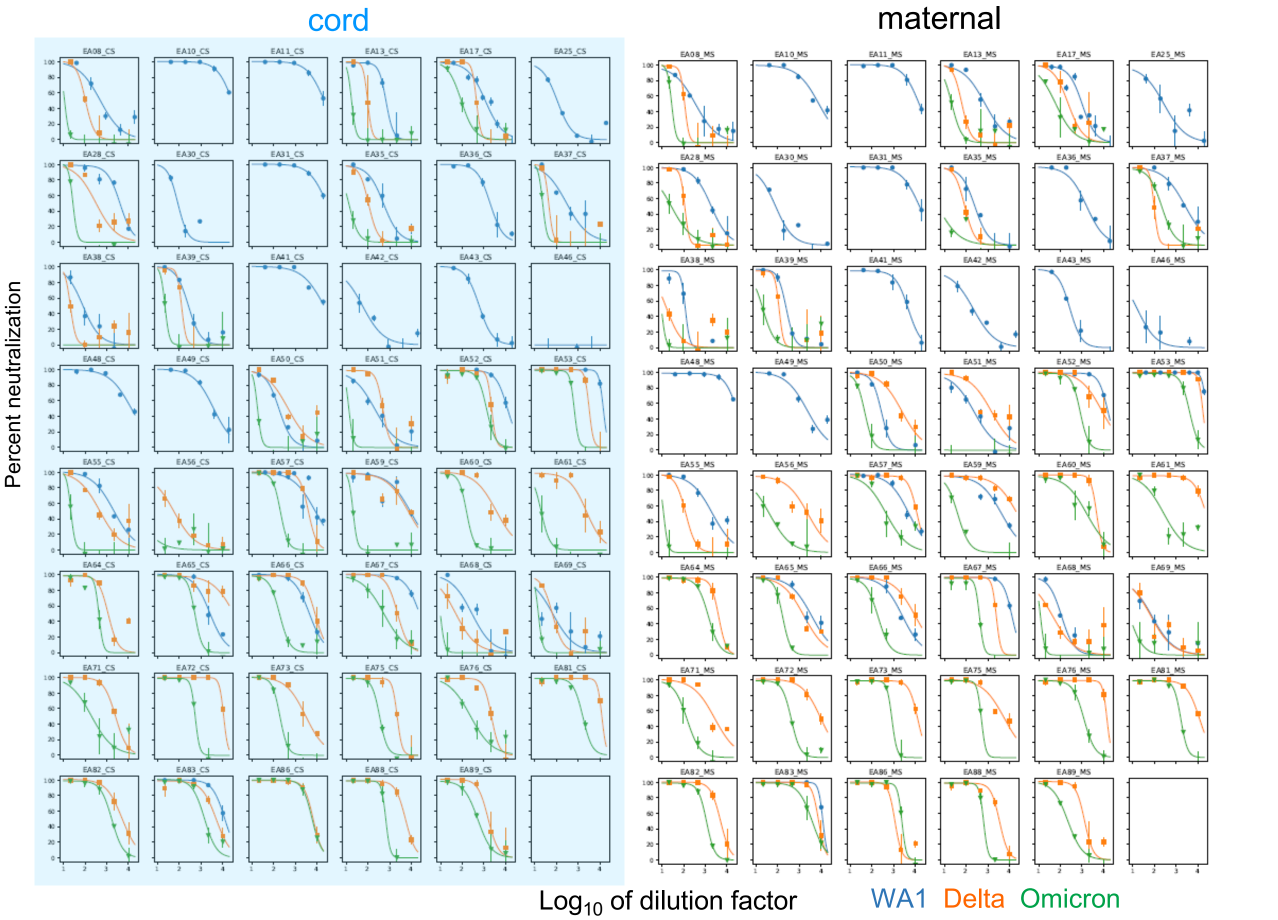


Supplemental Figure 2: FRNT50 were calculated from neutralization graphs generated in focus forming assays for each SARS-CoV-2 WT and clinical variants. Each graph shows the data for one individual sample.

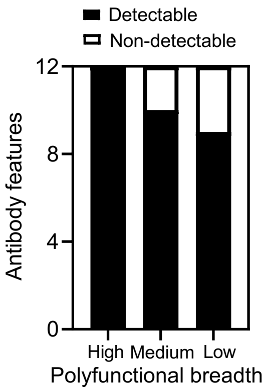


| Antibody features |
| --- |
| WA1 FRNT |
| Delta FRNT |
| Omicron FRNT |
| RBD ADNKA CD107a |
| RBD ADNKA IFNγ |
| RBD ADNKA TNFα |
| RBD ADCD |
| RBD ADCP |
| RBD CD16a |
| RBD CD32a |
| RBD CD32b |
| RBD FcRn |

Supplemental Figure 3: SARS-CoV-2 reactive polyfunctional breadth was calculated for each individual sample with all 12 features listed. In this cohort, individual responses fell into three main categories: those with high (90-100%), medium (80-90%) and low (<80%) proportion of functions detected.

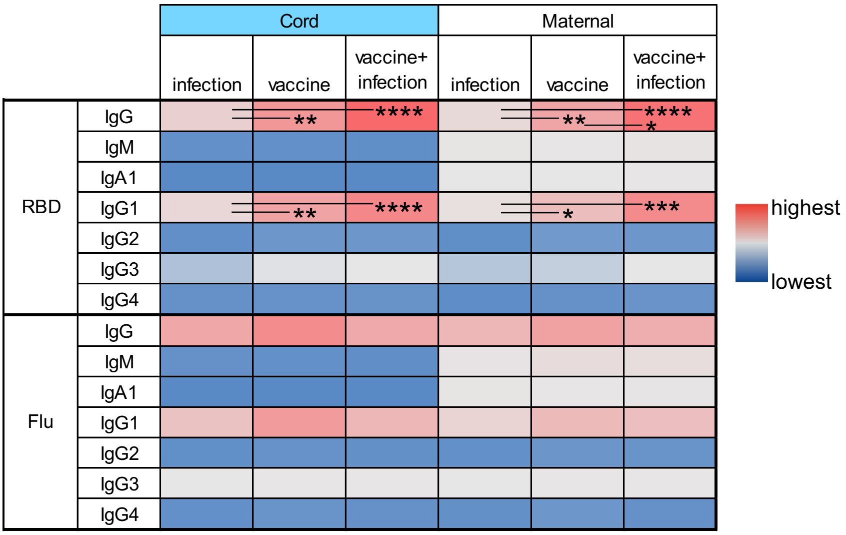


Supplemental Figure 4: No changes are observed in control influenza hemagglutinnin (Flu) compared to SARS-CoV-2 receptor binding domain (RBD) specific antibody levels in infant cord and maternal blood. The heatmap shows the median titers of antigen specific immunoglobulin isotypes (IgG, IgM, IgA1) and subclasses (IgG1, IgG2, IgG3, IgG4) within the three clinical groups from cord and maternal samples. P values are adjusted for maternal age and BMI using linear regression. * $p \leq 0.05$; ** $p \leq 0.01$; *** $p \leq 0.001$; **** $p \leq 0.0001$.

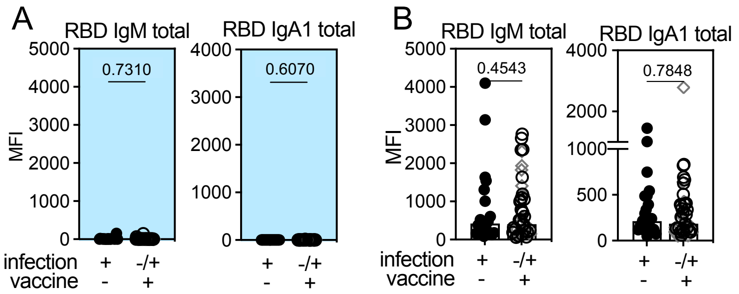


Supplemental Figure 5: No significant differences are observed in RBD IgM or IgA in maternal and infant cord blood with respect to immune exposure. Dot plots show the magnitude of RBD total IgM and IgA1 in (**A**) cord and (**B**) maternal samples. P values are adjusted for maternal age and BMI using linear regression.

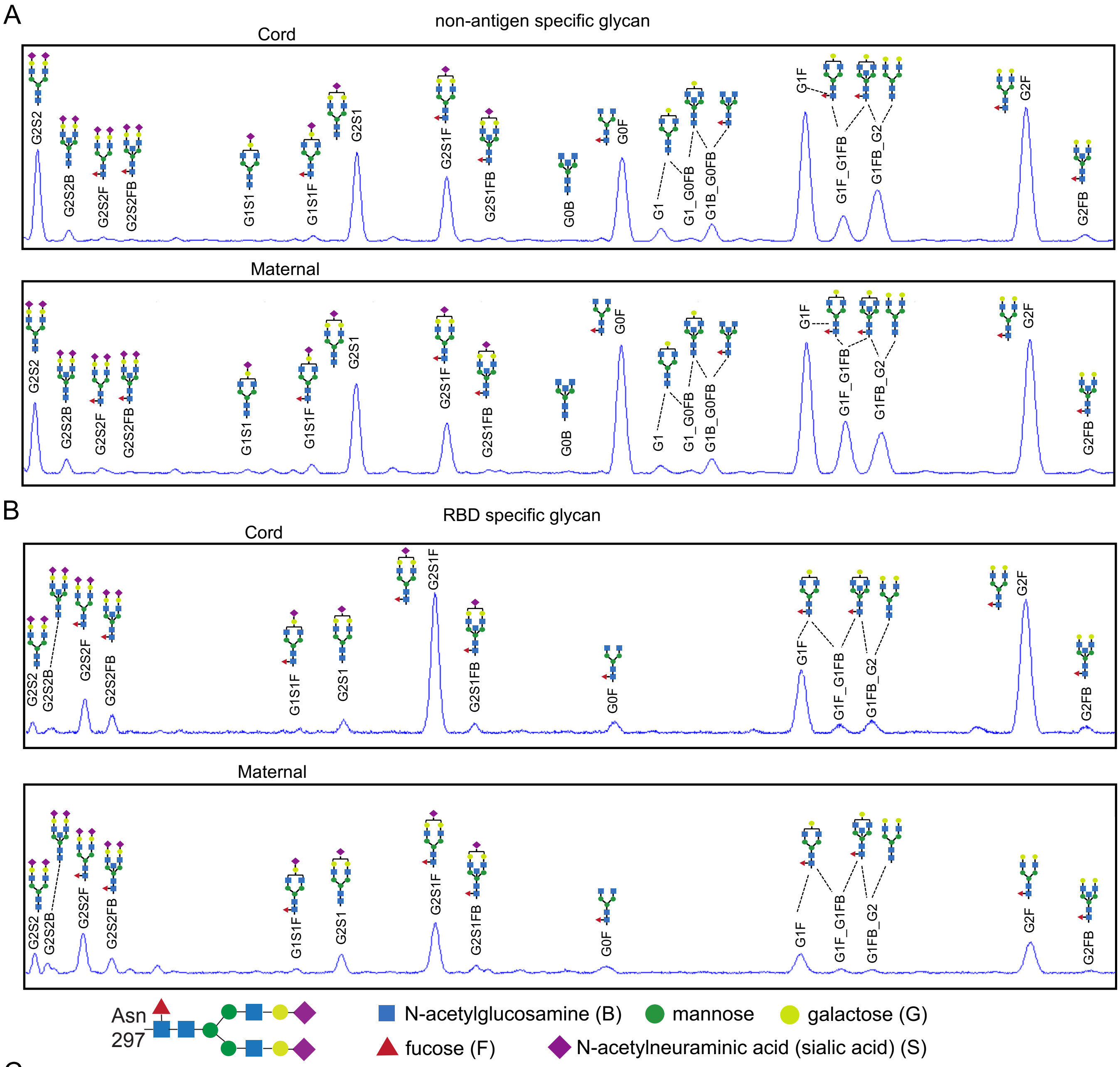


C

| Glycoform |
|---|
| total monogalactose (G1) = G1S1+ G1S1F+ G1+ G1_G0FB+ G1B_G0FB+ G1_2 +G1F+ G1F_G1FB+ G1FB_G2 |
| total digalactose (G2) = G2S2+ G2S2B+ G2S2F+ G2S2FB+ G2S1+ G2S1F+ G2S1FB+ G2F+ G2FB |
| total mono sialic acid (S1) = G1S1+ G1S1F+ G2S1+ G2S1F+ G2S1FB |
| total disialic acid (S2) = G2S2+ G2S2B+ G2S2F+ G2S2FB |
| total GlcNac (B) = G2S2B+ G2S2FB+ G2S1FB+ G0B+ G1B_G0FB+ G1FB_G2+ G2FB |
| total fucose (F) = G2S2F+ G2S2FB+ G1S1F+ G2S1F+ G2S1FB+ G0F+ G1F+ G1F_G1FB+ G1FB_G2+ G2F+ G2FB |

Supplemental Figure 6: Representative capillary electrophoresis chromatographs for one pair of maternal and infant cord samples show differences in relative RBD specific and non-antigen specific IgG glycosylation. Peaks are identified from bulk total non-antigen (**A**) and receptor binding domain (RBD) (**B**) specific IgG glycans captured and quantified by relative abundance. (**C**) The collective relative abundance of all individual glycoforms with fucose (F), sialic acid (S), galactose (G) and bisecting GlcNAc (B) are calculated.

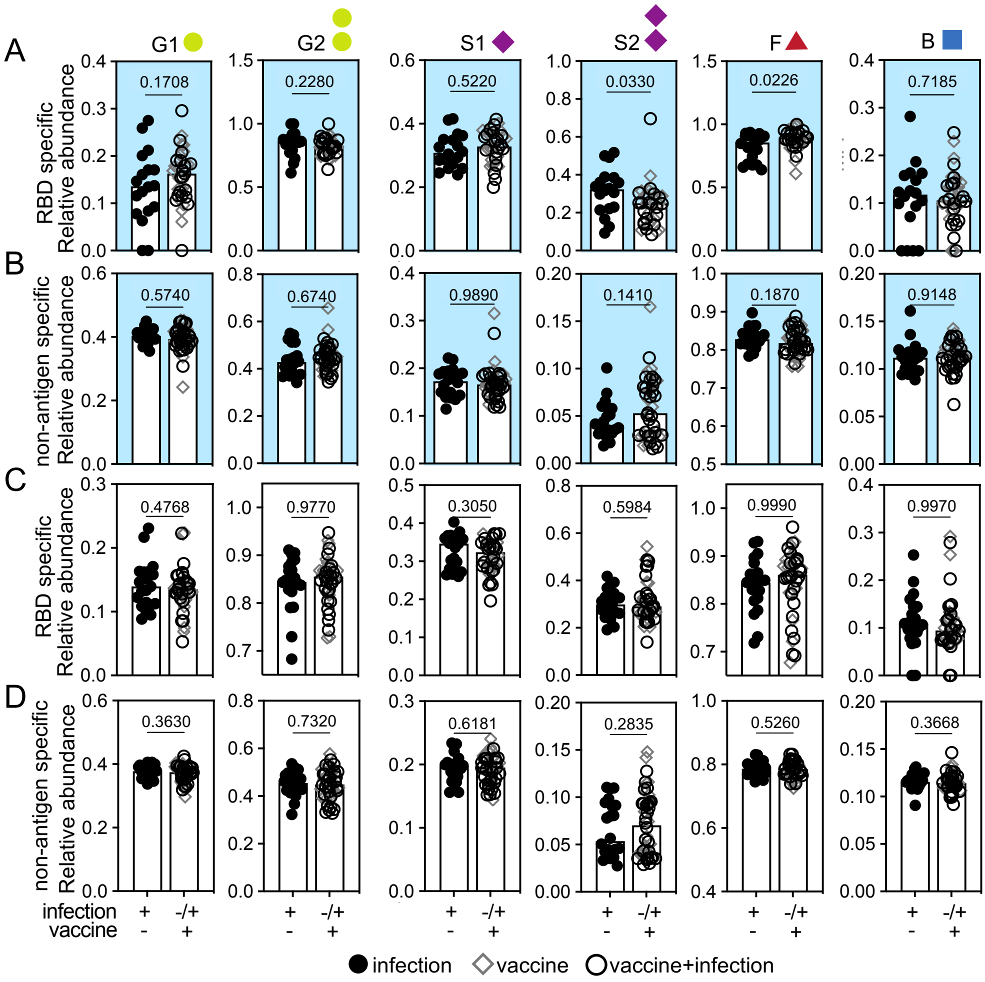


Supplemental Figure 7: Vaccination in pregnancy changes glycosylation of infant cord and not maternal RBD specific IgG. The relative abundance of glycoforms containing monogalactosylated (G1), digalactosylated (G2), monosialylated (S1), disialylated (S2) and bisecting N-acetylglucosamine (B) structures are depicted for RBD specific (**A** and **C**) and non-antigen specific (**B** and **D**) IgG from cord (**A** and **B**) and maternal (**C** and **D**) blood. For RBD specific IgG glycans, infection n=18, vaccine n=18, vaccine+infection n=19. For non-antigen specific IgG glycans, infection n=20, vaccine n=18, vaccine+infection n=27. Bars represent the median for each group. P values are adjusted for maternal age and BMI using linear regression.

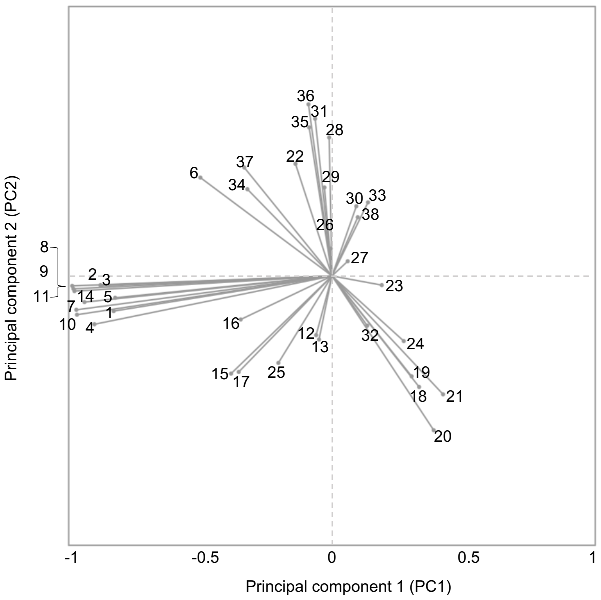


| Variable | RBD features | PC1 | PC2 |
|---|---|---|---|
| 1 | WA1 | -0.829365822 | -0.130694234 |
| 2 | CD107a | -0.879750329 | -0.03441598 |
| 3 | IFNγ | -0.875513728 | -0.036394076 |
| 4 | TNFα | -0.902394378 | -0.179475936 |
| 5 | ADCD | -0.82377986 | -0.081112146 |
| 6 | ADCP | -0.500077593 | 0.365078861 |
| 7 | CD16a | -0.971176365 | -0.125785108 |
| 8 | CD32a | -0.98683963 | -0.036924136 |
| 9 | CD32b | -0.982416837 | -0.047820267 |
| 10 | FcRN | -0.968775822 | -0.144040576 |
| 11 | IgG | -0.978406308 | -0.056341424 |
| 12 | IgA1 | -0.060366858 | -0.219808209 |
| 13 | IgM | -0.049312994 | -0.236050624 |
| 14 | IgG1 | -0.940278518 | -0.097271239 |
| 15 | IgG2 | -0.383402669 | -0.362721387 |
| 16 | IgG3 | -0.346759699 | -0.161701352 |
| 17 | IgG4 | -0.354224164 | -0.356439308 |
| 18 | G2S2 | 0.330937016 | -0.412227153 |
| 19 | G2S2B | 0.302646404 | -0.372773369 |
| 20 | G2S2F | 0.386347774 | -0.572894941 |
| 21 | G2S2FB | 0.421904424 | -0.439576598 |
| 22 | G1S1 | -0.138775776 | 0.416057965 |
| 23 | G1S1F | 0.189177582 | -0.034179648 |
| 24 | G2S1 | 0.272603426 | -0.241883427 |
| 25 | G2S1F | -0.203670579 | -0.323033316 |
| 26 | G0 | -0.005134646 | 0.101316486 |
| 27 | G1S1FB | 0.059655922 | 0.054464353 |
| 28 | G0B | -0.01090129 | 0.513848784 |
| 29 | G0F | -0.028533342 | 0.328413394 |
| 30 | G1 | 0.092375075 | 0.258601151 |
| 31 | G1_G0FB | -0.064162778 | 0.583706632 |
| 32 | G1B_G0FB | 0.133663612 | -0.184760757 |
| 33 | G1_2 | 0.136748855 | 0.27299171 |
| 34 | G1F | -0.32144356 | 0.321934859 |
| 35 | G1F_G1FB | -0.086844402 | 0.551447453 |
| 36 | G1FB_G2 | -0.089433477 | 0.637069014 |
| 37 | G2F | -0.333304865 | 0.402463557 |
| 38 | G2FB | 0.097743711 | 0.217831315 |

Supplemental Figure 8: Variability in all maternal and infant cord antibody data is captured most by SARS-CoV-2 antibody functions (PC1) and second most by RBD IgG glycans. PC scores of each of the 38 RBD specific antibody features in the loadings plot (left) are shown in the table (right).
